# Invasion of new adaptive zones retains telltale signs of directional selection at macroevolutionary scales in mammals

**DOI:** 10.64898/2025.12.21.695821

**Authors:** Fabio A. Machado, Anna Penna, Diogo Melo, Barbara A. Costa, Thiago M. G. Zahn, Ana C. Pavan, Arthur Porto, Harley Sebastião, Daniela M. Rossoni, Gabriel Marroig, Alex Hubbe

## Abstract

Directional selection is often viewed as a transient force in macroevolution, with its signal eroded over time by stabilizing and fluctuating selection. Yet, transitions into new adaptive zones are predicted to impose strong and sustained selective pressures that may leave a detectable signature even across deep timescales. We test this prediction by comparing the rates of multivariate skull morphological evolution required to traverse the boundaries between adaptive zones against genetic drift expectations. Our dataset includes 11,793 specimens spanning 231 species from 12 mammalian clades, each containing unique ecological transitions into new adaptive zones. Using a quantitative genetics framework, we estimated the phenotypic distances between ancestral and derived adaptive zones and contrasted them with null expectations under genetic drift. While a few adaptive zone invasions (e.g., marsupials and rodents) are consistent with drift, most exhibit substantially elevated rates of evolution. These results suggest that directional selection has recurrently shaped mammalian cranial evolution during major ecological shifts. We propose that adaptive zone transitions represent evolutionary contexts in which adaptation leaves a persistent macroevolutionary signal, challenging the prevailing view that long-term patterns are dominated by static forces.

## Introduction

Directional selection is widely regarded as the primary force driving macroevolutionary change. Yet, despite its central role in shaping phenotypic diversity, detecting its signature at large timescales remains remarkably difficult. This difficulty partly arises from 1) limitations of the tools used to identify directional selection and 2) the dynamics inherent to macroevolutionary data. Quantitative genetics methods typically infer evolutionary processes by contrasting empirical rates of phenotypic divergence with null expectations under genetic drift (Houle et al., 2017; Lande, 1985; Lynch, 1990; Turelli et al., 1988). However, macroevolutionary studies have shown a negative correlation between rates of evolution and the time period in which they are calculated, meaning that the longer the time span, the lower the rate of evolution is expected to be (Gingerich, 1983, 1987, 2001; Holstad et al., 2024). As a consequence, the signal of directional selection is expected to become increasingly obscured over time, making it difficult to distinguish it from changes generated by drift or stabilizing selection.

This pattern is consistent with the view that macroevolution is governed by adaptive landscape dynamics. Under this view, species remain strongly attracted to species-specific adaptive peaks until significant environmental shifts force changes in the distribution of peaks themselves (Hansen, 1997, 2024; Simpson, 1944; Uyeda et al., 2011). Because episodes of directional change are generally brief relative to the time spent evolving around adaptive optima (Eldredge & Gould, 1972; Simpson, 1944), rates of phenotypic evolution calculated over longer time intervals tend to decrease as they become dominated by periods of relative stasis, effectively erasing the signal of directional selection (Dzeverin, 2008; Lemos et al., 2001; Lynch, 1990; Machado et al., 2022; Porto et al., 2015; Uyeda et al., 2011). This effectively limits the usefulness of standard quantitative genetics approaches in macroevolutionary studies, especially for studying the signal of directional selection, leading to an emphasis on more phenomenological models of evolution that are less explicitly grounded in microevolutionary theory (Hansen, 1997).

While this emphasis on peak dynamics as the prevailing force has dominated the phylogenetic comparative methods literature (Hansen, 1997; Lynch, 1990; Uyeda et al., 2011), some studies did apply quantitative genetic methods to investigate macroevolutionary change directly. Most of these studies come from the biological anthropology/paleontology and primatology literature, in which rates of morphological evolution are compared with expectations derived from microevolutionary models. Studies of the divergence among human populations (Lynch, 1990; Roseman, 2004; Roseman & Weaver, 2007; Smith, 2011), between humans and Neanderthals (Weaver et al., 2007, 2008; Weaver & Stringer, 2015), during the evolution of the *Australopithecus*-*Homo* lineage (Schroeder et al., 2014), among higher apes (Hominidae; (Schroeder & von Cramon-Taubadel, 2017)), and across the primate order (Latrille et al., 2024; Machado et al., 2023) have reported rates of phenotypic divergence often consistent with drift and, in some cases, consistent indicative of directional selection

Investigations of this kind outside of primates have been limited and produced mixed results. Evaluations of morphological evolution in bats (Dzeverin, 2008) and marsupials (Lemos et al., 2001; Porto et al., 2015) found rates of evolution inferior to the expectation under genetic drift. A study on the evolution of Cingulata also found that rates were mostly slower than expected under drift, with a sole exception found in Glyptodons, a giant-sized fossil clade, for which the results were consistent with directional selection (Machado et al., 2022). Collectively, these findings suggest that the macroevolutionary signal of directional selection may persist under some circumstances, but also that its detectability varies across clades, traits, and evolutionary contexts. This uneven evidence indicates that the conditions under which directional selection remains detectable at macroevolutionary scales deserve broader comparative attention.

Here, we revisit this long-standing view that macroevolutionary rates are steadily consistent with stabilizing selection (Gingerich, 1983, 1987, 2001; Hansen, 1997; Lynch, 1990; Uyeda et al., 2011). Specifically, we test whether directional selection leaves a detectable macroevolutionary signature by quantifying the multivariate rates of cranial evolution across mammalian transitions into new adaptive zones, and by contrasting these rates with predictions from multivariate quantitative genetic models of drift. We focus on the mammalian skull because it is a semi-autonomous morphological system, highly informative about species ecologies,and therefore a likely target of natural selection (Machado, 2020; Marroig & Cheverud, 2004; Roseman, 2004). Adaptive zone transitions are particularly suitable for this test because they are expected to involve prolonged and intense periods of directional selection as lineages invade novel ecological regimes (Machado et al., 2022; Rossoni et al., 2017; Schroeder & von Cramon-Taubadel, 2017). By explicitly bracketing evolutionary transitions, we anticipate that we can diagnose events that required stronger directional selection, potentially leaving a significant signal even in comparative data.

## Materials and Methods

### Adaptive zone transitions

For more than 75 years, Simpson’s concept of “adaptive zones” has played a crucial role in our understanding of the evolutionary process over long timescales (Simpson, 1944, 1953). Under this model, an adaptive zone is a restricted region of the phenotypic space occupied by species with relatively similar form-function-fitness relationships (Polly, 2008; Simpson, 1953; Van Valen, 1971). Within-zone evolution is usually small in scale and does not involve major ecological changes. Evolution within a zone is thought to happen at exceptionally slow rates (*i*.*e*., bradytelic in Simpson’s terms), with lineages slowly following adaptive peaks within a restricted region of the morphospace. However, when opportunities arise (e.g., due to the invasion of new environments or extinction of competitors), lineages may experience faster rates of evolution (*i*.*e*., tachytelic) while invading new regions of the morphospace, leading to extreme evolutionary change associated with shifts in ecology and in the structure and distribution of adaptive peaks (Simpson, 1944, 1953; Uyeda et al., 2011).

Here, we applied the concept of adaptive zones to diagnose and define evolutionary transitions worth investigating. Because invasions of new adaptive zones are marked by increased rates of evolution, they are more likely to retain the signal of directional selection, even over longer timescales. Specifically, we use Polly (2008)’s formalization, which suggests that a phenotype is considered to inhabit a new adaptive zone if it a) evolved later; b) presents new ecological features; and c) falls outside the distribution of phenotypes that belong to the ancestral adaptive zone. To achieve this, within each clade, we classified species into two major ecological groups: one belonging to an ancestral group and the other to a recently evolved one. Species were assigned to a new adaptive zone if they differed in dietary or loco-motory ecology and are considered morphologically disjunct from the ancestral zone (see Supplementary material for more details).

Two general categories can be used to describe the relationship between adaptive zones. The first (henceforth, the “nested” type) corresponds to arrangements where lineages associated with the new adaptive zone are nested within a more inclusive clade of species belonging to the ancestral zone (*e*.*g*. Glyptodonts are nested within armadillos, therefore evolving from an armadillo-like ancestor; Figure 1A; Machado et al., 2022). In this case, we have direct access to phenotypes that represent the ancestral adaptive zone. The second category (the “analog” type) is based on identifying an extant analog for an extinct grade of species that preceded the invasion of the new adaptive zone (*e*.*g*., saber-toothed cats evolved from the *Pseudaelurus* grade of conical-shaped cats, functionally similar to Pantheridae; Figure 1B; (Simpson, 1944)). Using these analogs is essential because, in these cases, we did not have access to a direct representative of the potential ancestral morphologies.

**Figure 1:**
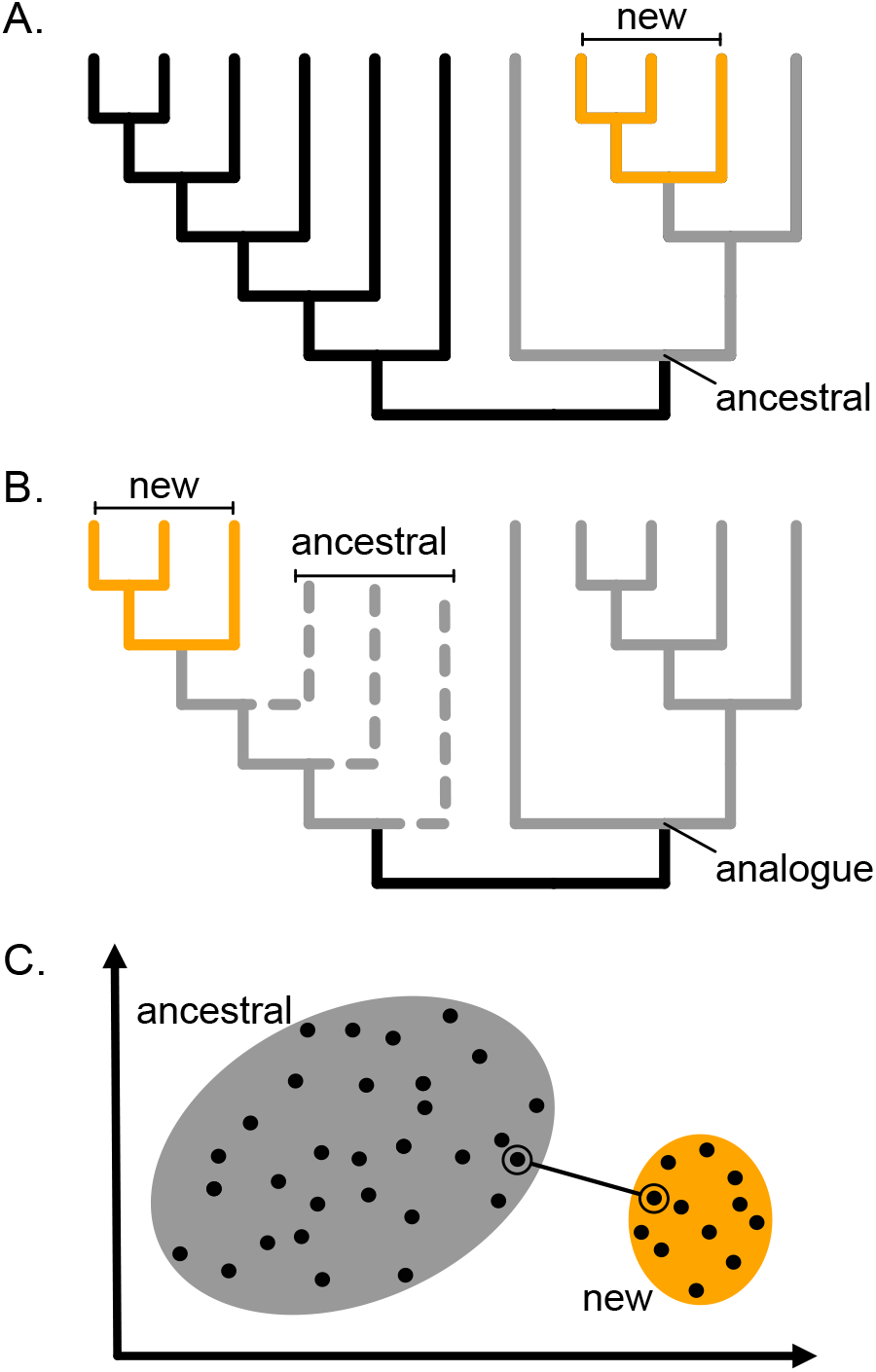
Examples of nested (A) and analog (B) adaptive zone transitions. Gray lineages are under the ancestral adaptive zone (or its analog), and orange lineages are under the new adaptive zone. Dotted lines represent extinct lineages. Ellipses on the hypothetical morphospace (C) represent both adaptive zones. The solid line connects the closest neighboring species used to model the transition among zones.

To model the invasion of a new adaptive zone, we identified, for each clade, the shortest distance between sampled phenotypes that leads to a transition between the ancestral and the new adaptive zone (Figure 1C, see below). The starting species is defined as the member of the ancestral adaptive zone that is morphologically closest to a member of the new zone, while the end species is the member of the new adaptive zone that is closest to the starting species. Even though these species are close in morphospace, they are generally not the closest phylogenetic relatives, as many modeled transitions involves different genera (Table 1).

**Table 1:**
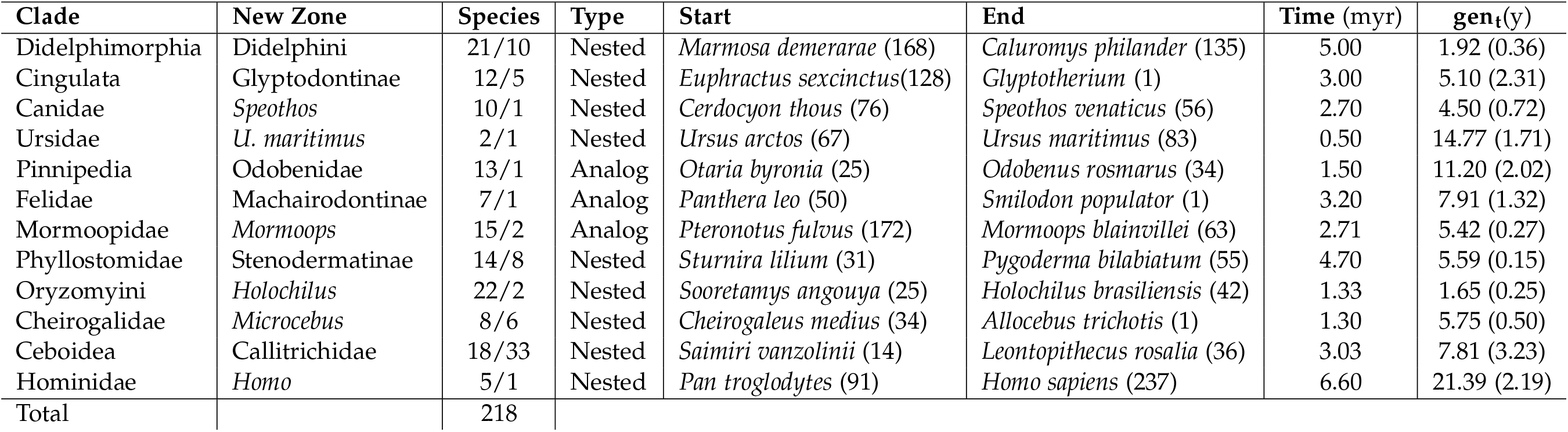
Adaptive zone transitions examined. **Clade**-Focal clades of analyses used to define ancestral adaptive zones. **New zone**-taxa thought to have colonized a new adaptive zone, **Species**-number of species in the ancestral zone and in the new zone, respectively. **Type**-Type of adaptive zone transition modeled. See text and figure 1A-B for full description. **Start** and **End**-Species chosen for the start and end of the adaptive zone transition. Numbers in parentheses are sample sizes. **Time** (myr) - Time in million years estimated for the transition according to phylogenetic and paleontological information. **gen**_**t**_(y) - Mean and standard deviation for the generation time in years for each clade.

We focused on 12 mammalian clades that exhibit between-lineage transitions consistent with the definition of adaptive-zone shifts used here (Table 1). These transitions were selected because the derived lineages differ from their putative ancestral zones not only in skull morphology, but also in ecological or functional attributes described independently of the present analyses. In Didelphimorphia, we examined the emergence of large-bodied South American opossums (Didelphini) from smaller didelphid ancestors. In Cingulata, we assessed the evolution of herbivorous, heavily armored glyptodonts (Glyptodontinae) from armadillo-like ancestors. In Canidae, we analyzed the evolution of the hypercarnivorous bush dog (*Speothos venaticus*) relative to more generalized South American canids. In Ursidae, we studied the divergence of the marine-specialized and hypercarnivorous polar bear (*Ursus maritimus*) from more generalist bears such as *U. arctos* and *U. americanus*. In Pinnipedia, we focused on the origin of mollusk-feeding walruses (Odobenidae) from more generalized otariid-like ancestors. In Felidae, we evaluated the shift from conical-toothed, panther-like felids to saber-toothed cats (Machairodontinae). In Mormoopidae, we examined the evolution of ghost-faced bats (*Mormoops*), which show distinctive craniofacial morphology and dietary specialization relative to more generalized insectivorous *Pteronotus*. In Phyllostomidae, we considered the emergence of specialized fruit-eating Stenodermatinae from insectivore-frugivore ancestors. In Oryzomyini, we traced the adaptation of *Holochilus* to semiaquatic habitats from terrestrial oryzomyine ancestors. Among primates, we focused on two independent cases of miniaturization: the evolution of mouse lemurs (*Microcebus*) within Cheirogaleidae and the evolution of marmosets and tamarins (Callitrichidae) within Ceboidea. We also examined the divergence of humans (*Homo sapiens*) from other apes within Hominidae, a transition associated with substantial craniofacial reorganization. A more thorough description, justification, and references for these transitions are provided in the Supplementary Material. From this point forward, we refer to each transition by the most inclusive taxon named above, and to the new adaptive zone by the less inclusive taxon (Table 1).

For the timing of each transition, we used information from phylogenetics and the fossil record to bracket the specific period of the invasion. For nested adaptive zones, this bracket typically lies between the origin of the first lineage in the new zone and the divergence time of a sister lineage still within the ancestral one. For the extant analog case, we used fossil-record information to bracket the origins of the last species belonging to the ancestral grade and the first species belonging to the new clade. See the supplementary file for specific information on each transition.

### Sample and Morphometrics

We evaluated the morphology of 231 species, comprising a total of 11,793 specimens (Table 1). Samples were obtained from 43 museums distributed worldwide (see acknowledgments). We primarily targeted adult specimens to minimize potential bias in estimating covariances arising from grouping across ontogenetic stages (Hubbe et al., 2023). Maturity was inferred based on complete dental eruption, fusion of the basicranial sutures, and overall morphology consistent with that of adults in each species (Hubbe et al., 2016; Machado et al., 2018). For Rodentia and Didelphimorphia, adult age classes were determined using dental age categories based on tooth eruption and wear (Hubbe et al., 2023). The age factor was statistically controlled for in these groups (see below). For Carnivora, we also included subadult specimens (skull size and shape consistent with adult morphology but without complete fusion of basicranial sutures), as their cranial measurements showed no significant differences from those of adults (Machado et al., 2018).

To quantify skull morphology, we calculated 33 standardized linear distances from 32 homologous anatomical landmarks, as described in (Machado et al., 2022). Landmarks were obtained using a Microscribe digitizer for most of our sample (de Oliveira et al., 2009; Hubbe et al., 2016; Machado et al., 2018; Pavan & Marroig, 2016; Porto et al., 2009; Rossoni et al., 2019), while the New World Monkey dataset was digitized with a Polhemus 3Draw digitizer (Marroig & Cheverud, 2001). Bilaterally symmetrical measurements were averaged within a specimen. Traits were set on the ratio (proportional) scale through log-transformation (Houle et al., 2011; Voje et al., 2023) following Holstad et al. (2024). To visually inspect species distribution across morphospaces, we performed principal component analysis (PCA) of the between-species covariance matrix.

For each species in our sample, we obtained a vector of average measurements, and the pairwise Euclidean distances between species in each clade were calculated. The shortest distance between species averages belonging to different adaptive zones was recorded as the distance needed for that transition, and the end-point species were selected for further analyses (Figure 1C, Table 1). Using other metrics (*e*.*g*., Mahalanobis distance) didn’t change our overall results.

### Rate test

To investigate how rapidly lineages would need to evolve to cross the phenotypic gaps between adaptive zones, we employed a quantitative genetics test based on the Lande Generalized Genetic Distance (LGGD) (Lande, 1979), calculated as

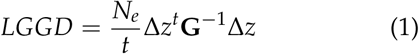

where *t* is the time of divergence in generations, *N*_*e*_ is the effective population size, Δ*z* is a vector representing the evolutionary transition for each trait, and **G** is the matrix of additive genetic variance and covariance among traits. The LGGD is thus a rate version of the Mahalanobis distance that uses quantitative genetics parameters to scale multivariate phenotypic change among lineages in units of standing genetic variation. So two transitions with the same phenotypic distance can have very different LGGD values if one occurred over a much shorter timescale or involved traits with less genetic variance available for evolution.

Solely under genetic drift, evolutionary change is expected to accumulate according to Lande’s equation of multivariate genetic drift (Lande, 1979):

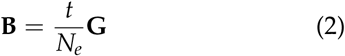

where **B** is the covariance matrix between evolutionary changes. This equation provides the null expectation against which we compared our observed distribution of LGGD. We used this method instead of expectations generated by parametric methods because the latter produce inadequate type-I error rates (Machado et al., 2022). In practice, LGGD values falling within the simulated drift interval can be interpreted as compatible with drift, whereas values substantially above or below indicate rates of evolution compatible with directional or stabilizing selection, respectively.

These methods depend on estimating **G**, which requires data from a population with known genetic relatedness (Falconer & Mackay, 1983). Because our samples come from museum collections with unknown relationships, we rely instead on the observation that phenotypic correlations — *i*.*e*. the sample correlation among traits — are a good approximation of the underlying genetic correlations (Cheverud, 1988), a fact thoroughly validated for mammalian skulls (Garcia et al., 2014; Hubbe et al., 2023; Marroig & Cheverud, 2010; Porto et al., 2015; Porto et al., 2009). Furthermore, phenotypic variance has been shown to scale linearly with additive genetic variance (Holstad et al., 2024). Together, these suggest that we can model **G** using phenotypic correlations and variances, *i*.*e*., the phenotypic co-variance matrix **P**.

We calculated **P** for each of the starting species (Table 1) by removing confounding effects (*e*.*g*., age, sex, geography) using a linear model approach (Marroig & Cheverud, 2001). Given that the sample sizes for some starting species were small, we also ran all analyses using the pooled-within-group covariance matrix between the start and end species. Because the results did not differ qualitatively when using these pooled matrices, we present only the results from a single-species matrix.

To address estimation uncertainty, we employed a Bayesian approach with a weak uninformative Wishart prior (Melo et al., 2015), drawing 1000 samples from the posterior distribution of Ps for each analysis. The use of uninformative priors has a similar effect as a shrinkage operation, which has been shown to reduce noise in the estimation of covariance matrices (Marroig et al., 2012). We then modeled **G** as

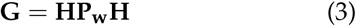

with **H** being a diagonal matrix with the square root of each trait’s *h*^2^ along the diagonal. This transformation assumes that phenotypic correlations approximate genetic correlations, while allowing the total amount of additive genetic variation to vary across traits and heritabilities. For *h*^2^, we drew values from a uniform distribution ranging from *h*^2^ = .3*™* .6, which is the range of values for the cranial traits explored here (Garcia et al., 2014; Marroig & Cheverud, 2010; Porto et al., 2015; Porto et al., 2009). The resulting distribution of *G*s thus incor-porates uncertainty in the estimate of genetic correlations (approximated by their phenotypic counterparts), phenotypic variances, and heritabilities.

To calculate the morphological change Δ*z* associ-ated with each evolutionary transition, we obtained a posterior distribution of the averages for the start and end species. Posterior parameters were obtained using a conjugate prior and drawing 1000 samples for each trait from the posterior distributions. Differences between posterior samples of the averages were then used to calculate 1000 Δ*z*s.

For *t*, we divided the divergence time available for each transition by the generation time (*gen*_*t*_). Cladewise averages and standard deviations of *gen*_*t*_ were obtained from Pacifici et al. (2013). We used these parameters to generate 1000 *gen*_*t*_ values using a normal distribution. For *N*_*e*_, we sampled 1000 values from a lognormal distribution with a mean of 11 and standard deviation of 1. The resulting *N*_*e*_ values ranging from 1,000–300,000, consistent with genomic estimates for multiple mammalian species (Brevet & Lartillot, 2021; Kuderna et al., 2023; Lorenzen et al., 2011; Wilder et al., 2023). Lastly, we plotted estimated LGGD against transition times to assess potential time dependence in the estimated rates.

## Results

As expected, across most focal clades, species assigned to new adaptive zones occupied regions of the morphospaces that were discontinuous from, or peripheral to, those occupied by species within the ancestral adaptive zones (Figure 2). The degree of separation, however, varied substantially among clades. Some transitions showed clear phenotypic discontinuities between ancestral and derived zones, including Glyptodontinae, Odobenidae, Machairodontinae, Mormoopidae, Ceboidea, and Hominidae (Figure 2). In these cases, the modeled transition crossed an evident gap between the two regions of morphospace. Other transitions were less sharply separated. Didelphimorphia, Oryzomyini, and Phyllostomidae showed new adaptive zones that extended from, or lay close to the edge of, the ancestral distribution (Figure 2), suggesting weaker phenotypic discontinuity between zones. Ursidae represents an ambiguous case: within-zone distances among the sampled bears are greater than the modeled between-zone distance involving polar bears. Because only three species were included, it is unclear whether the zones are adjacent or discontinuous.

**Figure 2:**
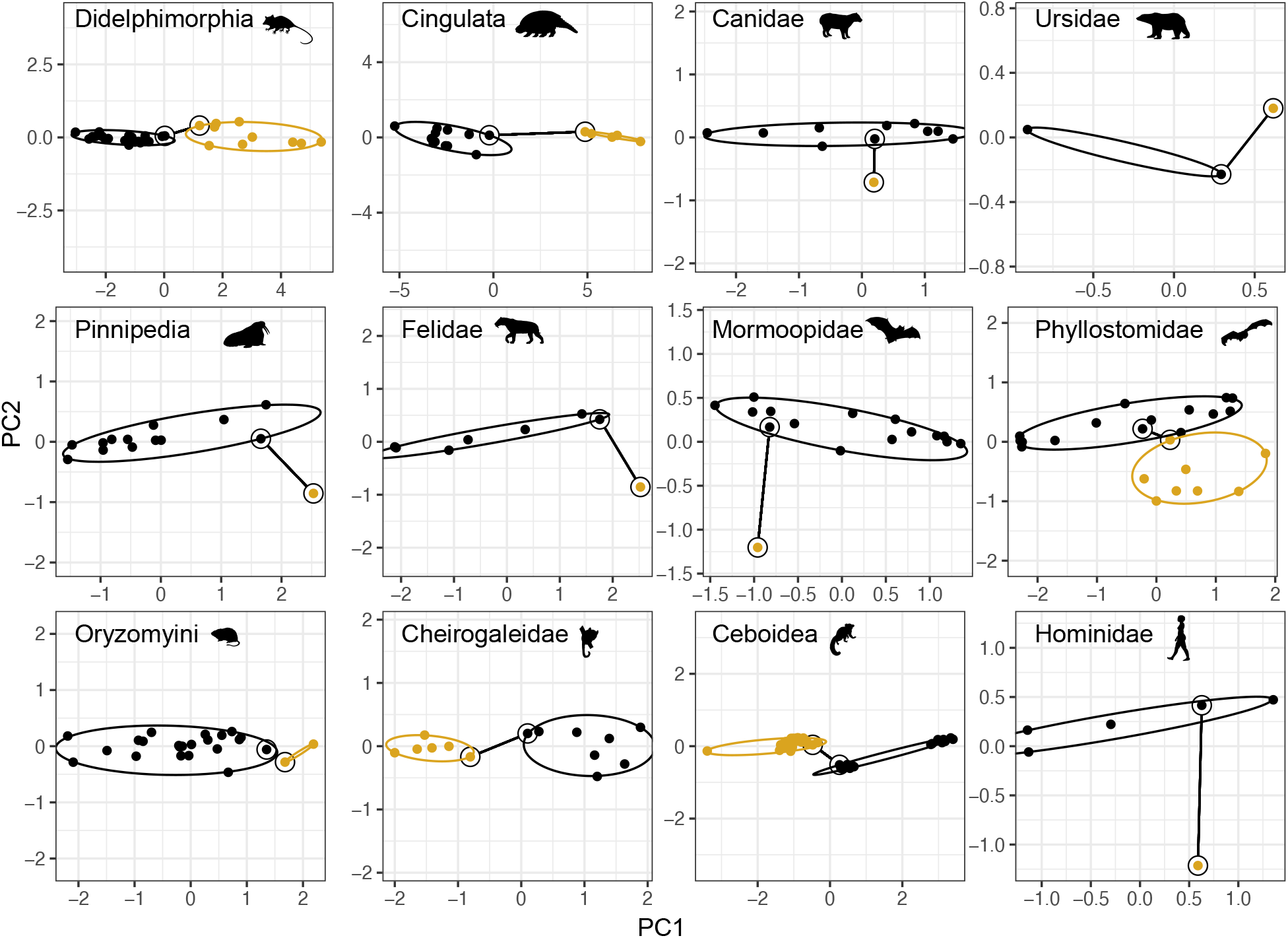
Distribution of the first two principal components of species averages for each focal clade. Species from the ancestral adaptive zone are represented in black, and the new adaptive zone is represented in gold. Curved solid lines represent the ellipsoidal envelope containing all species from a respective adaptive zone. Solid lines connect the pair of extreme species for each transition. Panels are in different scales.

The LGGD simulations under genetic drift produced lower and upper bounds of approximately 19 and 50 LGGD units (*D*^2^ *N*_*e*_*t*^*™*1^), respectively (Figure 3). Most modeled adaptive-zone transitions exceeded this interval, consistent with rates higher than expected under drift. The strongest signals were observed for Odobenidae and Glyptodontinae,whose LGGD distributions were clearly above the drift expectation. Several other transitions, including Felidae, Mormoopidae, Phyllostomidae, Ursidae, Ceboidea, and Hominidae, had median values, and often interquantile ranges, above the upper boundary expected for drift, although the magnitude varied among clades.

**Figure 3:**
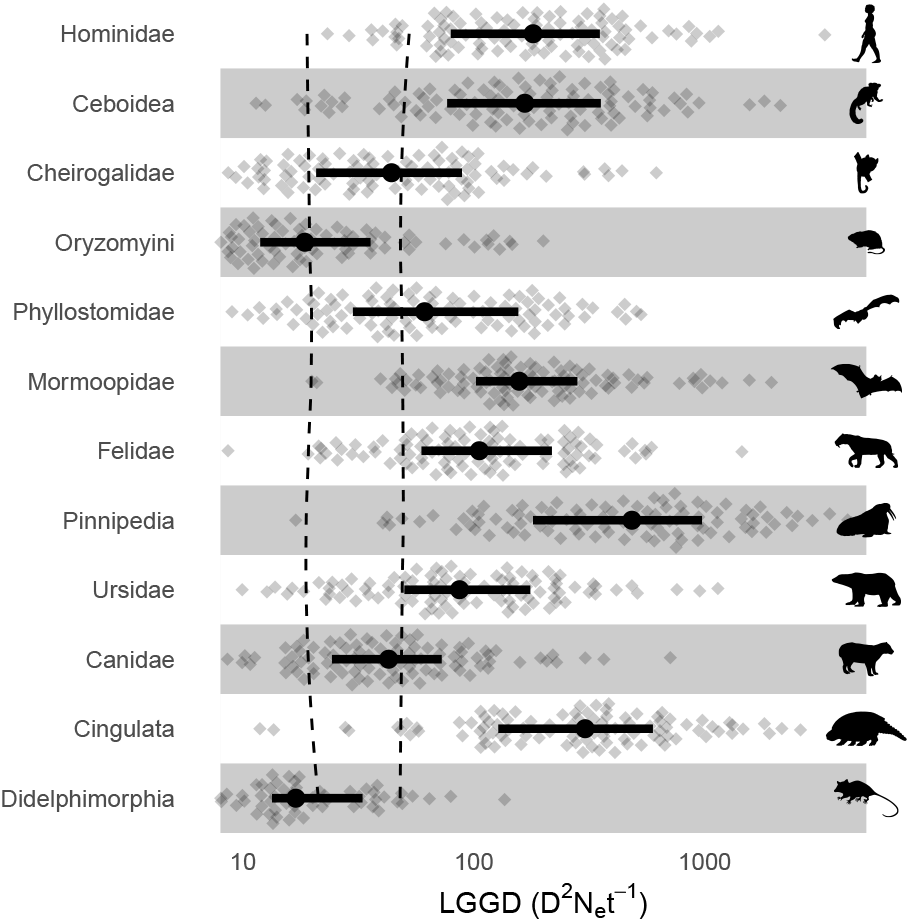
LGGD values for each adaptive zone transition. Diamonds represent each individual value of LGGD, dots are the medians and solid horizontal lines represent the interquantile ranges. Dashed vertical lines traversing all panels represent the lower and upper rates expected under drift estimated by simulation.

A smaller set of transitions showed weaker or no evidence of rates above drift expectations. Cheirogalidae and Canidae had median LGGD values near the upper bound of the drift expectation, suggesting borderline cases rather than strong departures from drift. Didelphimorphia and Oryzomyini fell below the expected drift interval, indicating that the modeled phenotypic transitions in these clades did not require elevated evolutionary rates. These low LGGD values also correspond to the weaker visual separation observed in the morphospace, particularly for Oryzomyini, in which the new adaptive zone lies near the edge of the ancestral distribution (Figure 2I). These results indicate that adaptive-zone transitions differ not only in whether they exceed drift expectations but also in the intensity of the evolutionary rates required to traverse the inferred phenotypic distances.

Contrary to the expectation that evolutionary rates tend to decline with increasing temporal interval, we found no consistent relationship between LGGD and time intervals (Figure 4). Pearson correlation coefficients calculated between each posterior LGGD sample and divergence time spanned both positive and negative values (*ρ* = *™*0.39 to 0.41), suggesting that longer transitions were not associated with lower rates of evolution or with a more muted signal of selection.

**Figure 4:**
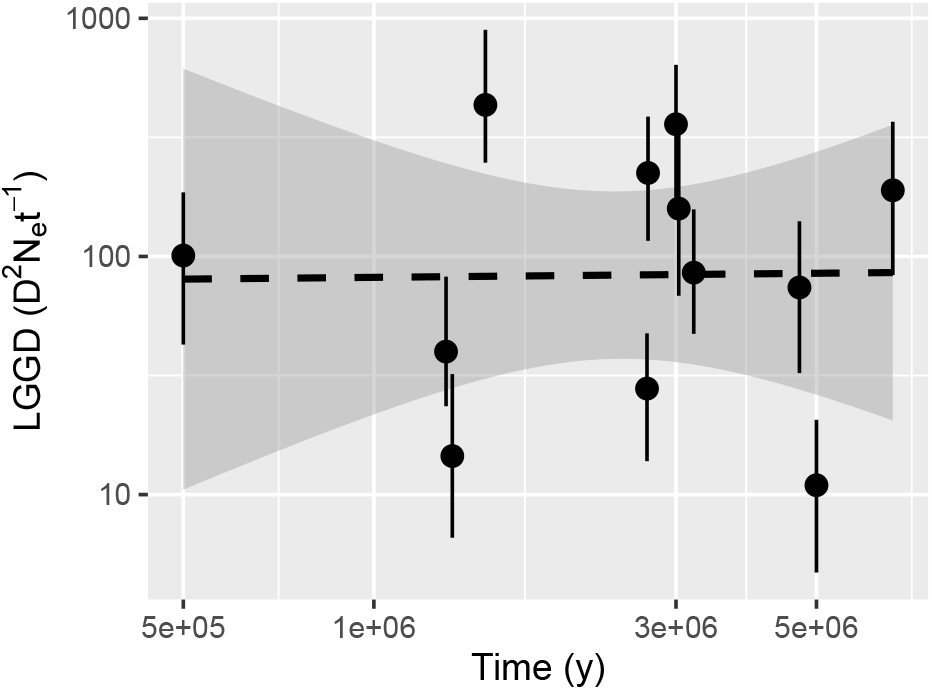
Relationship between rates of evolution, as measured by LGGD, and the timing of adaptive-zone transitions.

## Discussion

By focusing on evolutionary transitions more likely to involve intense directional selection, we recover here a pattern that contrasts with prevailing expectations from macroevolutionary theory. Evolutionary rates do not appear to decline with time (Fig. 4), nor were they constrained below the expectation under drift. Instead, most transitions exhibited rates at or above drift expectations (Fig. 3) despite spanning divergence timescales of 0.5–6.6 Myr (Table 1). These results suggest that the apparent “disappearance” of signals of directional selection at macroevolutionary scales might not be an inevitable consequence of temporal averaging. Rather, when we examine evolutionary change within biologically meaningful time intervals that bracket adaptive transitions, the signals of directional selection can remain detectable even over millions of years. More broadly, our find-ings support the idea that adaptive-zone invasions represent a class of evolutionary events in which adaptation and directional selection can leave persistent signals at macroevolutionary scales.

Among all tested transitions, those with the strongest signal of directional selection were walruses and glyptodonts. One possible explanation for this might be the direction of divergence between ancestral and new adaptive zones in these transitions. Both glyptodonts and walruses have skulls that are dorsoventrally expanded relative to their sister taxa (a phenomenon described for glyptodonts as “telescoping”; Farina and Vizcano, 2001). In mammals, size is typically the major axis of variation (Marroig & Cheverud, 2005), and thus the main direction of evolutionary potential (evolvability; Hansen and Houle, 2008), closely followed by independent changes in the relative size of the neurocranium and facial regions (Machado et al., 2018; Marroig & Cheverud, 2010; Rossoni et al., 2024). This might explain why adaptations along either size (*e*.*g*., Ceboidea, Cheirogalidae) or craniofacial length adjust-ments (*e*.*g*., Canidae, Ursidae, Phyllostomidae) did not require particularly high selective pressures – consequently displaying relatively lower LGGD values. However, since the evolution of walruses and glyptodonts involves variation along more constrained directions, greater intensity of selection is necessary to overcome the relative constraints imposed by patterns of trait covariation (Machado, 2020; Marroig & Cheverud, 2005). These results indicate that the detectability of directional selection depends on both the magnitude of ecological and morphological change and the alignment between evolutionary trajectories and the structure of available phenotypic variation.

The only clades falling below the expectation under drift are Didelphimorphia and Oryzomyini, which likely reflect clade-specific contingencies. In marsupials, size variation dominates evolutionary potential (Lemos et al., 2001; Porto et al., 2015), enabling rapid evolutionary responses. For rodents, however, the the presence of a highly versatile skull morphology (Maestri et al., 2017) might reduce the selective intensity required for ecomorphological change (Machado, 2020). Similar factors may account for the lower selection signals observed in Cheirogalidae and Canidae. For Cheirogalidae, differentiation between the new and ancestral zones occurred primarily along the size dimension, a well-known line of least resistance in primates (Marroig & Cheverud, 2005, 2010). On the other hand, while the selection pressure for a hypercarnivore diet in Canidae was mainly focused on reducing facial length to improve bite force (Machado, 2020), the facial region in this clade seems to have greater evolutionary flexibility (Machado et al., 2019; Machado et al., 2018), which might have facilitated adaptation in this direction.

Among bats, both groups showed rates above those expected under drift, with Phyllostomidae presenting lower values than Mormoopidae. This is a surprising result, since the former is routinely considered a classic example of adaptive radiation (Rossoni et al., 2017). Differences among groups, however, might relate to the same processes that led to the divergence of walruses and glyptodonts. The skull morphology of Stenodermatinae (obligate frugivore bats) is marked by an extreme reduction in facial length, which might have been facilitated by a more modular skull observed in Phyllostomids (Rossoni et al., 2024). Although a direct investigation of Mormoopidae skull modularity has not been conducted, it is unlikely that this group differs radically from established patterns for mammals or Chiroptera (Porto et al., 2009; Rossoni et al., 2019). In that case, the extreme inflection observed in the genus *Mormoops* is unlikely to align with any genetic line of least resistance, suggesting that it requires even greater selective pressure. The differences in the signal of selection among bat families further highlight the importance of context, rather than just morphological disparity and ecological shifts, in shaping selective dynamics (Schluter, 2024). In other words, ecological opportunity alone is insufficient to predict evolutionary rates. Instead, the interaction between ecological shifts and the structure of phenotypic integration may be crucial to determining the strength of directional selection required to reach a novel adaptive zone (Machado, 2020).

The results presented here are, of course, contingent on our modeling choices and the assumptions they entail. Some choices could inflate or underestimate the magnitude of the inferred rates, but most are unlikely to alter the central conclusion that adaptive-zone transitions often exceed drift expectations. For example, by defining adaptive zones from observed phenotypes, we necessarily treat sampled species as proxies for broader form-function-fitness relationships. Incomplete sampling, especially of fossil or unsampled extant intermediate forms, could therefore inflate distances between adaptive zones and, consequently, the inferred evolutionary rates. On the other hand, models of evolution that include pleiotropic effects on additional traits generally predict slower evolutionary responses than models that account only for focal traits (Hansen & Houle, 2008; Jiang & Pennell, 2025; Jiang & Zhang, 2020), implying that our estimates of the selection signal are in fact conservative, as they may underestimate the strength of selection required to produce the observed transitions. Therefore, while the exact magnitude of the inferred rates remains sensitive to modeling choices, the presence of elevated evolutionary rates across most adaptive-zone transitions appears robust to reasonable biological assumptions.

More caution may be warranted in our interpretation of transitions for which the new zone lies close to the edge of the ancestral distribution such as in Didelphimorphia and Oryzomyini. Because both groups exhibit highly conserved and presumably versatile cranial morphologies, assigning specific taxa to different adaptive zones may not be justified. In those cases, the lower rates of evolution found here could, in fact, be representative of within-zone bradytelic evolution (Simpson, 1944). By contrast, transitions such as Glyptodontinae, Odobenidae, Machairodontinae, and Hominidae involve clearer ecological and morphological discontinuities and are less likely to be explained solely by incomplete sampling of intermediate phenotypes. These cases, therefore, provide stronger evidence that adaptive-zone transitions can retain a detectable signal of directional selection.

Finally, this study is not intended as a methodological template for estimating the strength of selection during adaptive-zone transitions. Our goal is more limited: to test whether directional selection can leave detectable signatures in macroevolutionary data when transitions are defined and temporally bracketed using independent comparative, phylogenetic, and paleontological information. We therefore selected cases that were especially likely to involve sustained adaptation into new adaptive zones. As a result, our findings show that such signals can persist under favorable conditions, but they should not be used to infer how frequently directional selection remains detectable across macroevolutionary datasets more generally.

We note, however, that previous studies using comparative quantitative-genetic frameworks have reached similar conclusions (Latrille et al., 2024; Machado et al., 2022; Machado et al., 2023; Schroeder & Ackermann, 2017; Schroeder et al., 2014). Together with these studies, our results challenge the assumption that directional selection is necessarily obscured at macroevolutionary timescales. They instead suggest that adaptive-zone transitions may provide especially favorable contexts for detecting how microevolutionary processes contribute to large-scale phenotypic diversification.

Future work should move beyond asking whether signals of directional selection can persist and instead test how often, and under what ecological contexts, they shape the tempo of evolution. Achieving this will require models of phenotypic evolution that more fully integrate comparative phylogenetic methods, quantitative-genetic frameworks, and fossil data.

## Data and code availability statement

Necessary data and code to replicate all analyses are available at https://github.com/MachadoFA/AdapZones.

## Author contributions

FAM and AH developed the research question. AH collected data on Cingulata. BAC collected data on rodents. FAM and TMGZ collected data on carnivores. GM and AP collected data on primates. ACP and DMR collected data on bats. AP and HS collected data on Didelphidae. FAM and DM developed the analytical and computational analyses. FAM wrote the first draft, and all authors contributed to later drafts and discussion.

## Acknowledgments

We thank the curatorial staff of the following institutions for allowing access to their collection: American Museum of Natural History (AMNH), British Museum of Natural History (BMNH), *Coleção de Chiroptera de São José do Rio Preto* (DZSJRP), Duke Primate Center (DUPC), *Facultad de Ciencias Naturales y Museo de La Plata* (FCNyM), Field Museum (FMNH), *Faculdad de Ciencias de la Universidad de la Republica* (FC-UDELAR), Florida Museum of Natural History (UF), *Instituto de Pesquisas Amazonicas* (INPA), *Instituto de Pesquisas Científicas e Tecnológicas do Estado do Amapá* (IEPA), Kentucky University (KU), Los Angeles County Museum (LACM), *Naturhistorisches Museum, Wien* (NMW), *Museo Argentino de Ciencias Naturales Bernardino Rivadavia* (MACN), *Museo de Historia Natural de la Universidad Nacional Mayor de San Marcos* (MUSM), *Museo Municipal de Colonia* (MMC), *Museo Nacional de Historia Natural* (MNHNUY), *Museu de Anatomia Humana da Universidade Federal de São Paulo* (MAH-UNIFESP), *Museu de Anatomia Humana Professor Alfonso Bovero* (MAHPAB), *Museu de Ciências Naturais da Fundação Zoobotânica do Rio Grande do Sul* (MCNRGS), *Museu de Ciências Naturais da PUC-MINAS* (MCN), *Museu de Zoologia da Universidade de São Paulo* (MZUSP), *Museu Nacional do Rio de Janeiro* (MNRJ), *Museu Paraense Emílio Goeldi* (MPEG), *Museum d’Histoire Naturelle, Geneve* (MHNG), *Museum für Naturkunde* (MfN, ZMB), *Muséum National d’Histoire Naturelle* (MNHN-FR), Museum of Comparative Zoology (MCZ), Museum of Vertebrate Zoology (MVZ), *Naturhistorisches Museum, Bern* (NHM), Royal Belgian Institute for the Natural Sciences (RBINS), Royal Museum for Central Africa (RMCA), Royal Ontario Museum (ROM), *Senckenberg Naturmuseum Frankfurt* (SMF), Smithsonian Institute (USNM), Texas Tech University (TTU), *Universidade Federal da Minas Gerais* (UFMG), *Universidade Federal da Paraíba* (UFPB), *Universidade Federal do Pernambuco* (UFPE), *Universidade Federal do Piauí* (UFPI), University Museum of Zoology, Cambridge (UMZC), *Zoologisches Forschungsmuseum Alexander Koenig* (ZFMK). We also acknowledge M. Teixeira, P. Auricchio, and V. Tavares for access to specimens under their care. We thank Josef Uyeda for his insights into a previous version of the manuscript and Leonardo Borges for his assistance with Figure 1.

## Supplementary material

### Justification for transition choices

In Didelphimorphia, we examined the emergence of large-bodied South American opossums in Didelphini from smaller-bodied didelphid ancestors. Larger didelphimorphia experience significantly diverse ecological and biomechanic pressures, leading to more morphological diversification than what is observed in small-bodied species (Abreu & Astúa, 2025; Amador & Giannini, 2016; Astúa, 2009; Silva-Neto et al., 2024).

In Cingulata, we assessed the evolution of Glyptodontinae from armadillo-like ancestors. Glyptodonts represent a distinctive herbivorous, heavily armored radiation within Cingulata, which shows significant functional differences from extant armadillos (Cuadrelli et al., 2020; Machado et al., 2022; Vizcaíno et al., 2011).

Among Carnivora, we investigated four transitions: within Canidae, Ursidae, Pinnipedia and Felidae. In Canidae, we analyzed the evolution of the bush dog *Speothos venaticus*. These South American canids possess multiple cranial and mandibular specializations associated with hypercarnivory, in contrast to the more generalist South American canids (Beisiegel & Zuercher, 2005; Machado, 2020; Perini et al., 2010; Ruiz et al., 2023; Van Valkenburgh, 1991). In Ursidae, we studied the divergence of the polar bear *Ursus maritimus* from more generalist brown and black bears. This transition was accompanied by strong ecological, dietary, and physiological specialization for Arctic environments and marine mammal prey, rather than the more omnivorous diet seen in Ursinae (Liu et al., 2014; Polly, 2024; Slater et al., 2010). In Pinnipedia, we focused on the evolution of walruses (Odobenidae). Walruses have a highly modified tusked and suction-feeding skull, diverging greatly from more primitive members of the group, which were generalized piscivores closely resembling otariids(Boisville et al., 2024; Hocking et al., 2021; Horikawa, 1994; Repenning, 1976). In Felidae, we evaluated the transition from conical-toothed felids to saber-toothed Machairodontinae. Saber-toothed cats are characterized by elongate upper canines, reflecting a disparate strategy for prey dispatching than that seen in conical-toothed felids (Chatar et al., 2022; Emerson & Radinsky, 1980; Martin, 1980; Piras et al., 2018; Tamagnini et al., 2023).

Among bats, we analyzed two groups: Mormoopidae and Phyllostomidae. In Mormoopidae, we examined the evolution of ghost-faced bats *Mormoops*. Mormoops exhibits one of the most extreme morphologies in bats, with a pronounced dorsal inflection of the palate, a feature likely associated with a specific echolocation strategy and dietary specialization on lepidopterans(Gilley et al., 2025; A. Pavan et al., 2019; Rolfe et al., 2014). In Phyllostomidae, we considered the emergence of Stenodermatinae, a diverse radiation of specialized fruit-eating bats whose craniodental morphology is associated with the processing of soft and hard fruits, from insectivore-frugivore ancestors (Dumont et al., 2012; Rossoni et al., 2017).

In Oryzomyini, we traced the evolution of the marsh rats *Holochilus* from terrestrial oryzomyine ancestors. This genus is highly adapted to aquatic habitat use, possessing webbed feet and strong craniodental specializations for plant feeding (Soto et al., 2026; Tulli & Carrizo, 2024).

Among primates, we focused on two independent cases of miniaturization: the evolution of mouse lemurs *Microcebus* within Cheirogaleidae, including the smallest living primate species (Andrews et al., 2020; Masters et al., 2014); and the evolution of small-bodied callitrichids within Ceboidea, a group whose reduced body size and associated life-history traits distinguish them from other New World monkeys (Andrews et al., 2020). Finally, in Hominidae, we examined the divergence of *Homo sapiens* from other apes, a transition associated with marked craniofacial reorganization and previously inferred directional selection along the *Homo* lineage (Schroeder & Ackermann, 2017; Schroeder & von Cramon-Taubadel, 2017).

For each transition, we used phylogenetic or fossil evidence to bracket the time interval over which the adaptive-zone transition plausibly occurred. For Didelphini, we used the interval between the divergence of Didelphini from the remaining Didelphidae and the divergence of *Metachirus* from the remaining Didelphini (Silva-Neto et al., 2024). For Glyptodontinae, we used the interval between the molecularly estimated split of glyptodonts from other armadillos and the occurrence of early specialized glyptodonts such as *Propalaehoplophorus* (Cuadrelli et al., 2020; Delsuc et al., 2016; Mitchell et al., 2016; Vizcaíno et al., 2011). For *Speothos*, we used the timing of South American canid colonization after the formation of the Isthmus of Panama and the Great American Biotic Interchange (Perini et al., 2010). For *Ursus maritimus*, we used molecular estimates of divergence from brown bears (Hailer et al., 2012; Liu et al., 2014). For Odobenidae, we used the interval between late stem odobenids with generalized feeding morphologies, such as *Pontolis*, and the origin of derived Odobeninae (Boessenecker et al., 2023). For Machairodontinae, we used the interval between the last *Pseudaelurus*-grade conical-toothed felids and the origin of saber-toothed machairodontines (Jiangzuo et al., 2023; Paijmans et al., 2017). For *Mormoops*, we used the interval between the split of *Mormoops* from *Pteronotus* and the oldest fossil mormoopids (Morgan et al., 2019; A. C. Pavan & Marroig, 2017). For Stenodermatinae, we used the interval between the split of Stenodermatinae from the ancestral phyllostomid pool and the origin of crown Stenodermatinae (Rojas et al., 2016). For *Holochilus*, we used the interval between the origin of the ancestral lineage leading to *Holochilus*+*Pseudoryzomys* and the cladogenetic event separating these genera (Percequillo et al., 2021). For *Microcebus*, we used the interval between the origin of *Allocebus* and the emergence of *Microcebus* (Wisniewski et al., 2022). For Callitrichidae, although the split from other extant New World monkeys is older, fossil evidence places taxa such as *Lagonimico* and *Aotus dindensis* near the stem of callitrichine evolution and suggests body sizes compatible with larger-bodied New World monkeys; we therefore used the interval between these fossil stem taxa and the origin of crown Callitrichidae to bracket the invasion of the miniaturized callitrichid adaptive zone (Fleagle, 2013; Kay, 1994; Wisniewski et al., 2022). For *Homo*, we used the divergence time from chimpanzees estimated by Schroeder and von Cramon-Taubadel (2017).

